# Metabolic Control of Glycosylation Forms for Establishing Glycan-Dependent Protein Interaction Networks

**DOI:** 10.1101/2024.10.30.621210

**Authors:** Xingyu Liu, Li Yi, Zongtao Lin, Siyu Chen, Shunyang Wang, Ying Sheng, Carlito B. Lebrilla, Benjamin A. Garcia, Yixuan Xie

## Abstract

Protein-protein interactions (PPIs) provide essential insights into the complex molecular mechanisms and signaling pathways within cells that regulate development and disease-related phenotypes. However, for membrane proteins, the impact of various forms of glycosylation has often been overlooked in PPI studies. In this study, we introduce a novel approach, glycan-dependent affinity purification followed by mass spectrometry (GAP-MS), to assess variations in PPIs for any glycoprotein of interest under different glycosylation conditions. As a proof of principle, we selected four glycoproteins—BSG, CD44, EGFR, and SLC3A2—as baits to compare their co-purified partners across five metabolically controlled glycan conditions. The findings demonstrate the capability of GAP-MS to identify PPIs influenced by altered glycosylation states, establishing a foundation for systematically exploring the Glycan-Dependent Protein Interactome (GDPI) for other glycoproteins of interest.

## INTRODUCTION

Following the success of the Human Genome Project, researchers have achieved significant advancements in the Human Proteome Project over the past decade (Smith et al. 2021). As of today, over 90% of the human proteome has been uncovered. The next challenge lies in functional characterization of each identified protein and proteoform. Given that proteins do not operate in isolation within living organisms, a crucial aspect for investigating protein functions is through their protein-protein interactions (PPIs) (Armingol et al. 2021). Mass spectrometry (MS) has been a pivotal technology in the high-throughput approaches to identify PPIs. Common approaches often coupled with MS to study PPIs include affinity purification, proximity-based labeling, and cross-linking. Each method has its optimal application in detecting PPIs (Low et al. 2021; Smits and Vermeulen 2016). For instance, proximity-based labeling methods like BioID are suitable for detecting transient interactions, whereas crosslinking methods can be employed in cases where genetic editing is not feasible. On the other hand, affinity purification coupled with quantitative mass spectrometry analysis (AP-MS) is a classic and highly practical approach for many types of PPI studies (Dunham, Mullin, and Gingras 2012). This method is also highly flexible, easily combinable with other techniques, or streamlined for systematically constructing PPI networks. A milestone in constructing proteome-wide PPI networks was established by Huttlin and colleagues using the AP-MS approach (Huttlin et al. 2021; Huttlin et al. 2017; Huttlin et al. 2015). While all MS-based methods for mapping PPIs and their variations have their advantages and limitations, a significant challenge for almost all of them is resolving the changes caused by protein modifications. Most proteins undergo co-translational or post-translational modifications, generating different proteoforms and strongly influencing the parent protein function or activity (Lin and Caroll 2018). These protein modifications introduce a new dimension due to their inherent complexity, and the impact of the modification is inevitably overlooked during the construction of PPI networks (Wang, Osgood, and Chatterjee 2022).

A key example of this complexity is glycosylation, and the glycosylated proteins are crucial components of the highly interactive outer cell membrane layer known as the glycocalyx.(Varki et al. 2022) The biological functions of the glycocalyx can be significantly affected by variations in glycosylation. For example, glycosylation of the integrin beta 1 (ITGB1) is essential for protein expression and heterodimeric formation (Isaji et al. 2009). Glycans are also crucial for disintegrin and metalloprotease 10 (ADAM10) processing and resistance to proteolysis (Escrevente et al. 2008). Recent work has put forth different approaches to monitor glycoprotein interactions. For example, we and MacMillan groups introduced proximity-based techniques, POSE, POFE, and GlycoMap, to modify the sialylated and fucosylated glycoprotein and to profile their local microenvironments (Xie et al. 2024; Li et al. 2019; Meyer et al. 2022). Sun *et al*. also presented a system that relied on labeling galactose oxidase (GAO) and enables the interrogation of pertinent glycoprotein counter-receptors on the surface (Sun, Suttapitugsakul, and Wu 2021). In addition, we and others demonstrated the detection of different chemical crosslinking between sialic acids and their interacting proteins (Li et al. 2023; Xie et al. 2021). These techniques enabled the capture of the direct interaction between glycan and proteins. Glycosylation also induces conformational changes in proteins, thus affect their PPIs besides those directly mediated by glycans (Shental-Bechor and Levy 2008). The treatment of tunicamycin or PNGase F allows the discrimination of protein-protein and protein-glycan interactions (Joeh et al. 2020). Classic mutagenesis of the amino acid that carries glycosylation followed by the AP-MS approach can also help to resolve the glycan-dependant protein-protein interactions; however, the complete loss of glycosylation barely happens during biological processes, and, instead, glycoproteins mostly undergo the alternation of glycan types they carry (glycoform change), which can affect the protein state and its binding partners and thus exhibit different biological functions. Therefore, a platform that enables the deciphering of protein-protein interactions with different glycoforms is urgently desired and has great potential to impact glycoscience.

Recently, we employed a set of glycan modifiers for metabolic manipulation of glycan phenotype in cultured cells (Lebrilla et al. 2024). In this system, we took advantage of a human colorectal carcinoma cell line, HCT116, which bears the fucosylation deficiency caused by the mutation in GDP-mannose-4,6-dehydratase (GMDS) (Moriwaki, Shinzaki, and Miyoshi 2011). Treating the HCT116 cells with different glycan modifiers, including fucose, 3fluorinated sialic acid (3-F-Sia), and Kifunensine (Kif), could generate cells with different global glycan states. This approach produced five major glycan phenotypes: sialylated (S), neutral (Neu), fucosylated (F), sialofucosylated (FS), and high mannose (HM) types. Under each type, cells carry a dominant form of glycosylation. We reasoned that this system for controlling glycan phenotypes could be integrated with AP-MS to identify changes in PPIs resulting from specific types of glycosylations for theoritically any glycoprotein of interest (**Figure 1**). We name this new technique glycan-dependent affinity purification mass spectrometry (GAP-MS) analysis, a comprehensive platform that allows us to explore the glycoprotein interactome under different glycan phenotypes and provides novel insights into interactions affected by the various glycosylation forms (**Figure 1B-C**). We selected four bait glycoproteins, including BASI (Basigin), EGFR (epidermal growth factor receptor), CD44, and SLC3A2 (amino acid transporter heavy chain SLC3A2), spanning 156 high confident interactions as the initial study (**Figure 1D-E**). While many of these interactions have been covered by existing databases, nearly 45% are newly identified. This validates the methodology and demonstrates the sensitivity of our workflow for monitoring glycoprotein interactions. Importantly, we found that most of the interactions (131 out of 156) were involved in constructing the glycan-dependent protein interactome (GDPI), exhibiting strong preferences or dislikes across different glycan phenotypes. These results can be visualized on our website (www.glycointeractome.org). Finally, we performed mutagenesis on the glycosylation sites of BSG followed by AP-MS analysis, and compared the results with the BSG data from GAP-MS. The outcomes showed rare overlaps in the affected PPIs. Our results highlight the utility of the GAP-MS technique for revealing unprecedented insights into the glycoprotein interaction network.

**Figure 1.**
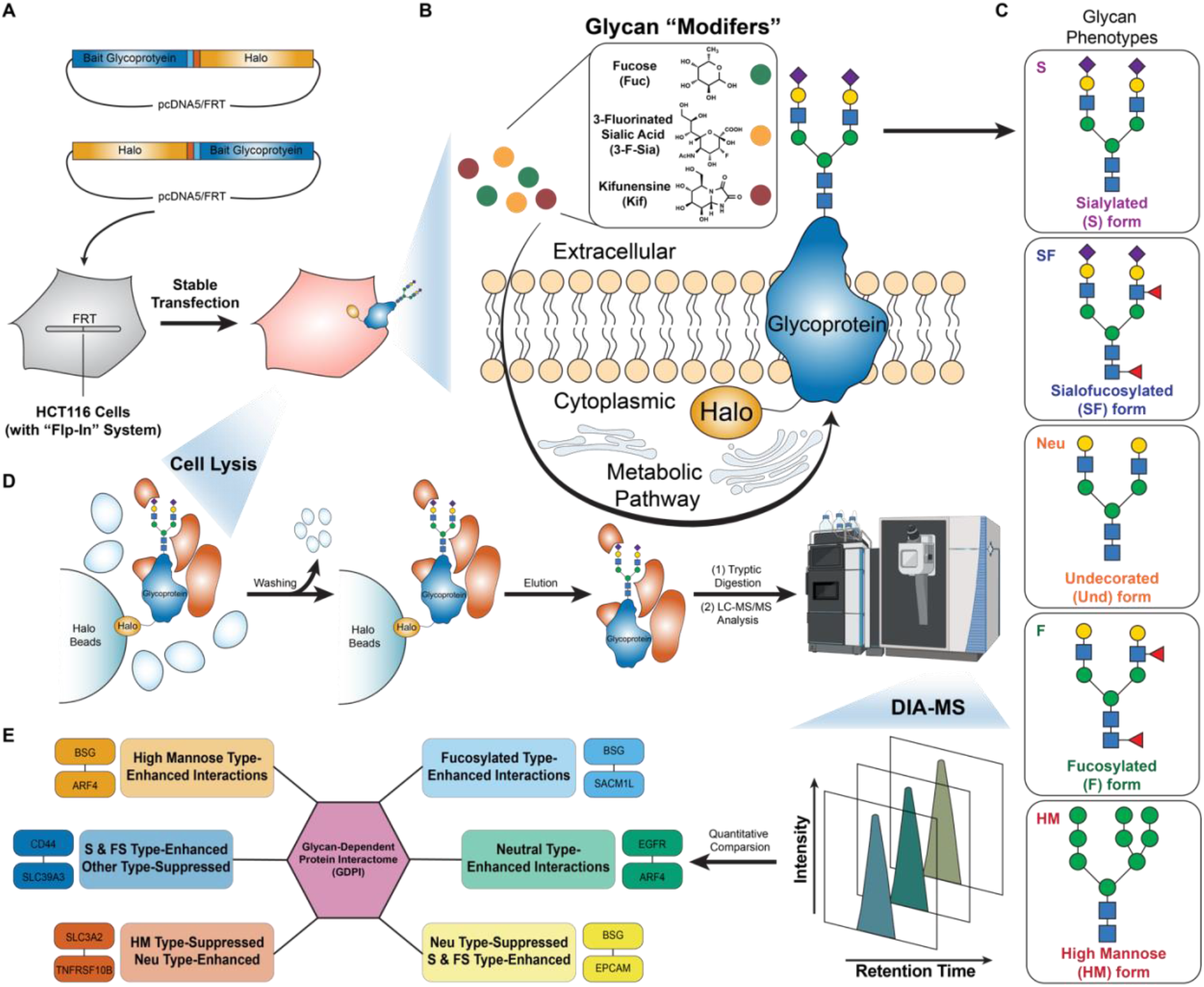
Schematic diagram of GAP-MS. GAP-MS employed glycan modifiers (e.g., fucose, 3-fluorinated sialic acid, and Kifunensine) to manipulate glycan phenotypes in HCT116 cells, generating five distinct phenotypes: sialylated (S), neutral (Neu), fucosylated (F), sialofucosylated (FS), and high mannose (HM). Integrating this approach with HaloTag-based affinity purification mass spectrometry (AP-MS) and data-independent acquisition (DIA) allows for the exploration of Glycan-Dependent Protein Interactome (GDPI).

## RESULTS

### Establishing glycan phenotypes on HCT116 membrane

We first created different glycan phenotypes on the membrane of the human colorectal carcinoma cell line, HCT116. We chose the HCT116 cell line over other cells because of its unique feature that the mutation of dehydratase GMDS leads to failure synthesizing GDP-fucose from GDP-mannose *via* the *De Novo* pathway and causes the lack of fucose source and fucosylation deficiency (Moriwaki, Shinzaki, and Miyoshi 2011). Importantly, the fucose can still be incorporated *via* the *Salvage* pathway by treating cells with exogenous fucose monosaccharide. We took advantage of these features and combined glyco-inhibitors of sialic acid and mannosidase to generate the system with five different glycan phenotypes on the cell membrane. Specifically, HCT116 cells are dominated by sialylated glycans without any treatment, while the glycans can be replaced with the SF type with external fucose added. With the treatment of a sialic acid inhibitor, 3-F-Sia, the Neu type can be generated, and with a combination of 3-F-Sia and Fuc, the F type is initiated. Lastly, the Kif treatment disturbs the mannosidase activity, the downstream biopathway is hindered, and Man9 and Man10 type of HM glycans are retained.

To examine whether the glycan profile could be altered with glycan modifiers, we perform glycomic analysis to profile the *N*-glycans under five conditions. We extracted the cell membrane, released the *N*-glycan using PNGase F, and mapped the *N*-glycans profile (**Supplementary Data 1A**). As shown in **Figure 2A-B**, the natural HCT116 showed a complete deficiency of fucosylation and was dominated by sialylated glycans (S form). At the same time, the treatment of fucose yielded cells with sialofucosylated glycans (SF form), while the co-treatment of 3-F-Sia and fucose converted the glycan with fucosylation (F form). The sole treatment of 3-F-Sia produced more than 55% neutral glycans without sialic acid or fucose (Neu form). Notably, four major glycan phenotypes were observed, with variations in the relative abundance of high mannose glycans, attributed to differences in glycan ionization efficiency. Lastly, the mannosidase inhibitor generated cells with more than 95% of high mannose glycans (HM form).

**Figure 2.**
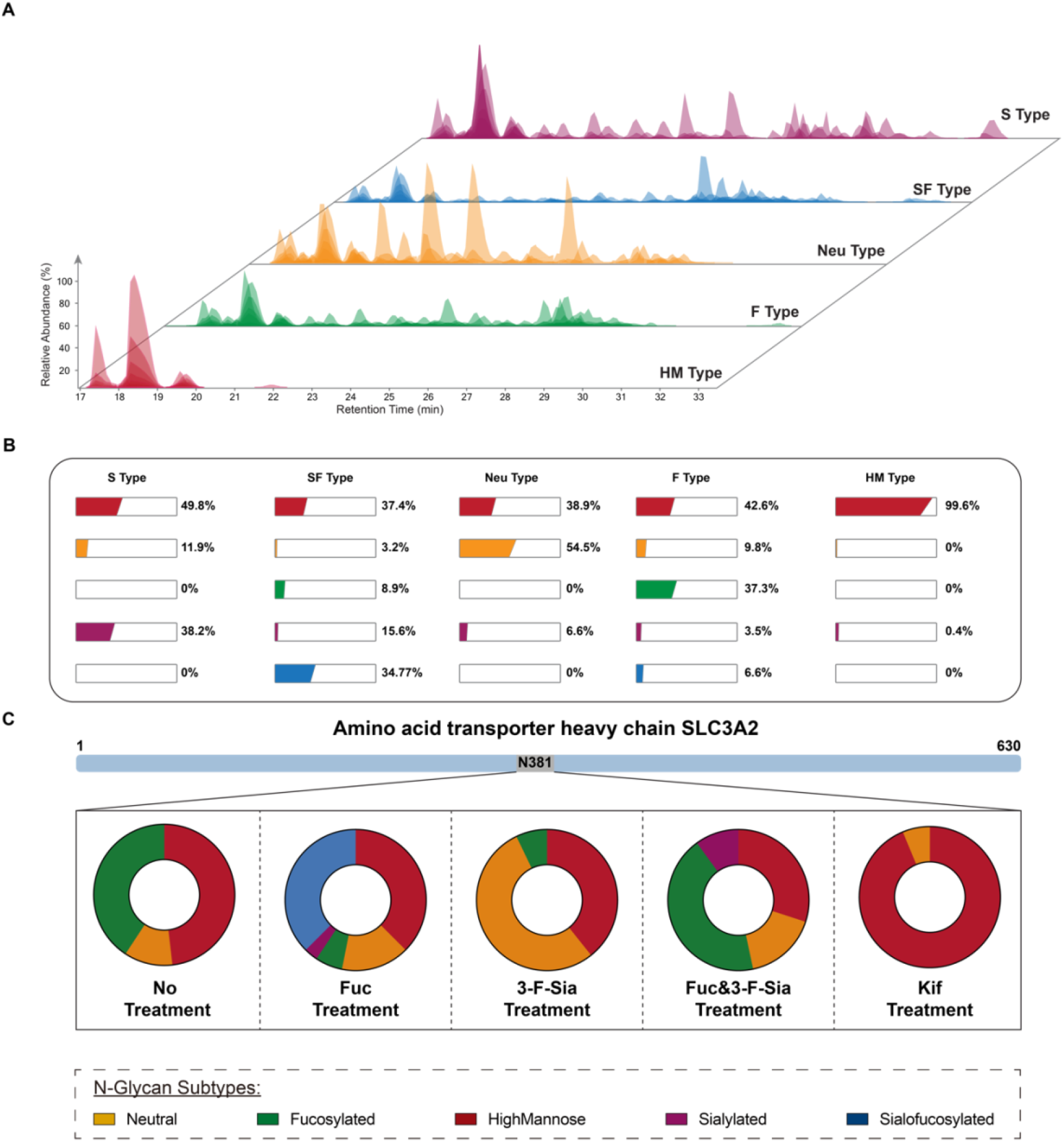
Glycomics and glycoproteomics monitored the glycan profiles after treating HCT116 cells with different modifiers. **(A)** Chromatogram of N-glycome profiles under five glycan conditions. **(B)** The relative abundance of N-glycans from glycomic analysis confirmed the generation of S, SF, F, and Neu phenotypes, with an additional HM phenotype induced by a mannosidase inhibitor. **(C)** The phenotypes of the bait glycoproteins were further confirmed through glycoproteomic analysis. As an example of the SLC3A2 bait glycoprotein, the glycans were altered with different treatments.

For a deeper investigation of the resulting glycan phenotypes at the glycopeptide level, we applied the glycan information from global glycan release as a focused library search and elucidated the information about both glycan and peptides with high confidence. The HILIC cartridge was employed to enrich glycopeptides specifically, and MS analysis enabled sensitive detection of glycopeptides and the site-specific glycoproteomic information (Li et al. 2020). As a result, we successfully identified 682 N-glycosites on 440 cell membrane glycoproteins, giving rise to over 2800 nonredundant glycopeptides in total (**Supplementary Data 1B**). The correlation between glycoprotein and different glycan types showed that glycan phenotypes are successfully produced for different glycoproteins at different glycosites. Collectively, the glycomic and glycoproteomic results both demonstrated the five major glycan phenotypes can be efficiently generated using the glycan modifiers.

Considering that the interacting protein level will be quantified and compared under these five conditions for the following AP-MS experiment, we want to ensure that the different glycan modifier treatments do not lead to a significant change in proteome profile. Hence, we employed the proteomic analysis under five conditions and evaluated protein abundance changes. Compared to the control condition (S type), we observed minimal changes in protein levels after treatment with glycan modifiers (**Figure S1** and **Supplementary Data 2**), which is consistent with previous observations from Caco-2 and A549 cell lines (Zhou et al. 2021). To be noted, FucFSia and Kif treatments generated more protein level floating compared to all the others, while there was no apparent enrichment of membrane proteins in significantly changed proteome under any condition. Taken together, our results emphasize the glycan modifiers only lead to the glycan expression level change instead of the whole proteome change.

### Producing bait glycoprotein with glycan phenotype expression on the cell membrane

As a proof of concept, we initiated the identification of glycan-dependent PPIs using GAP-MS with several examples of glycoproteins of interest as baits. As guided by the identified hub proteins from previous studies, we selected four bait proteins in this initial study, including CD44, BASI, EGFR, and SLC3A2 (Xie et al. 2021). Importantly, these baits are common glycoproteins found in various cells and with relatively high abundance in native environments, while glycans are crucial in regulating their diverse biological functions as observed previously (Varki 2017; Xie et al. 2020). Thus, we could map a comprehensive subnetwork of PPIs on the cell membrane from the interactions of these four proteins before systematically applying GAP-MS to a larger collection of bait proteins.

To perform affinity purification, we overexpressed selected glycoproteins of interest with HaloTag® in HCT116 cells. To minimize the interference of the tag to the glycosylation sites of the bait proteins, which often fall in the extracellular regions of transmembrane proteins, the HaloTag® was fused to the cytoplasmic termini of each bait. HaloTag® is a versatile tag that can also be used for fluorescent imaging of the tagged protein (Liu et al. 2024; Liu et al. 2020). This feature allows us to check the overexpressed bait proteins localized to the cytoplasmic membrane (**Figure S2A**). We employed the Flp-In™ technology to generate HCT116 cells that stably express Halo-tagged proteins. All stable-expression cell lines were derived from the same clone of HCT116 that carries the flippase recognition target (FRT) recombinant site, thus each bait glycoprotein was integrated at the same locus in the genome. The treatments of glycan modifiers to the stable-expression cells were the only experimental procedure before cells were collected as materials for affinity purification. This system minimized sample variations by avoiding changes caused by different glycan modifiers.

To confirm the glycans on the over-expressed bait proteins were sufficiently regulated by the glycan modifiers, we elucidated the site-specific glycopeptide information of the four proteins in the stable bait-expressed cell lines. Consistent with the results above, bait glycoproteins with different N-glycoforms were predominantly generated under five conditions. As an example shown in **Figure 2C**, the SLC3A2 bait owed over 50% of sialylated glycan natively, and the treatment of Fuc and 3-F-Sia converted those glycans into sialofucosylated and undercoated glycans, respectively. With the treatment of both Fuc and 3-F-Sia, cells were present with fucosylated glycans, while the high-mannose glycans were dominant in cells with the addition of Kif. The data confirmed the glycans on the bait glycoproteins were exquisitely controlled in our system.

### Deciphering glycoprotein interaction network

*Next*, we identified and quantified the membrane proteins purified with the bait protein using the data-independent acquisition (DIA) proteomics workflow, which provides better quantitative measurements and is beneficial for comparing the strength of interactions under different glycan conditions. Integrating the quantitative results with the Significance Analysis of INTeractome (SAINT) analysis, we could, with high confidence, identify the main interactors enriched by bait proteins compared to HaloTag mock control (**Supplementary Data 3A and B**). Using a SAINT score cutoff of 0.90, we identified a total of 85 interacting proteins and 156 interactions from GAP-MS analysis of the 4 bait proteins. This global PPI network is shown in **Figure 3A**. As the bait proteins are known to be interacting with each other in existing PPI databases, we first checked the module only containing BSG, CD44, EGFR, and SLC3A2 in our new network (**Figure S2B**). We could capture the known interactions between our baits including BSG-EGFR, BSG-CD44, EGFR-CD44, and BSG-SLC3A2. Examining the overlap between our network and the established database, over 46% of interactions (72 out of 156) from our platform were found on the STRING database with high confidence scores, 43 out of 156 interactions were recorded in the BioGRID database, and 66 interactions were newly identified by GAP-MS (Mering et al. 2003; Stark et al. 2006) (**Figure S2C-D**).

**Figure 3.**
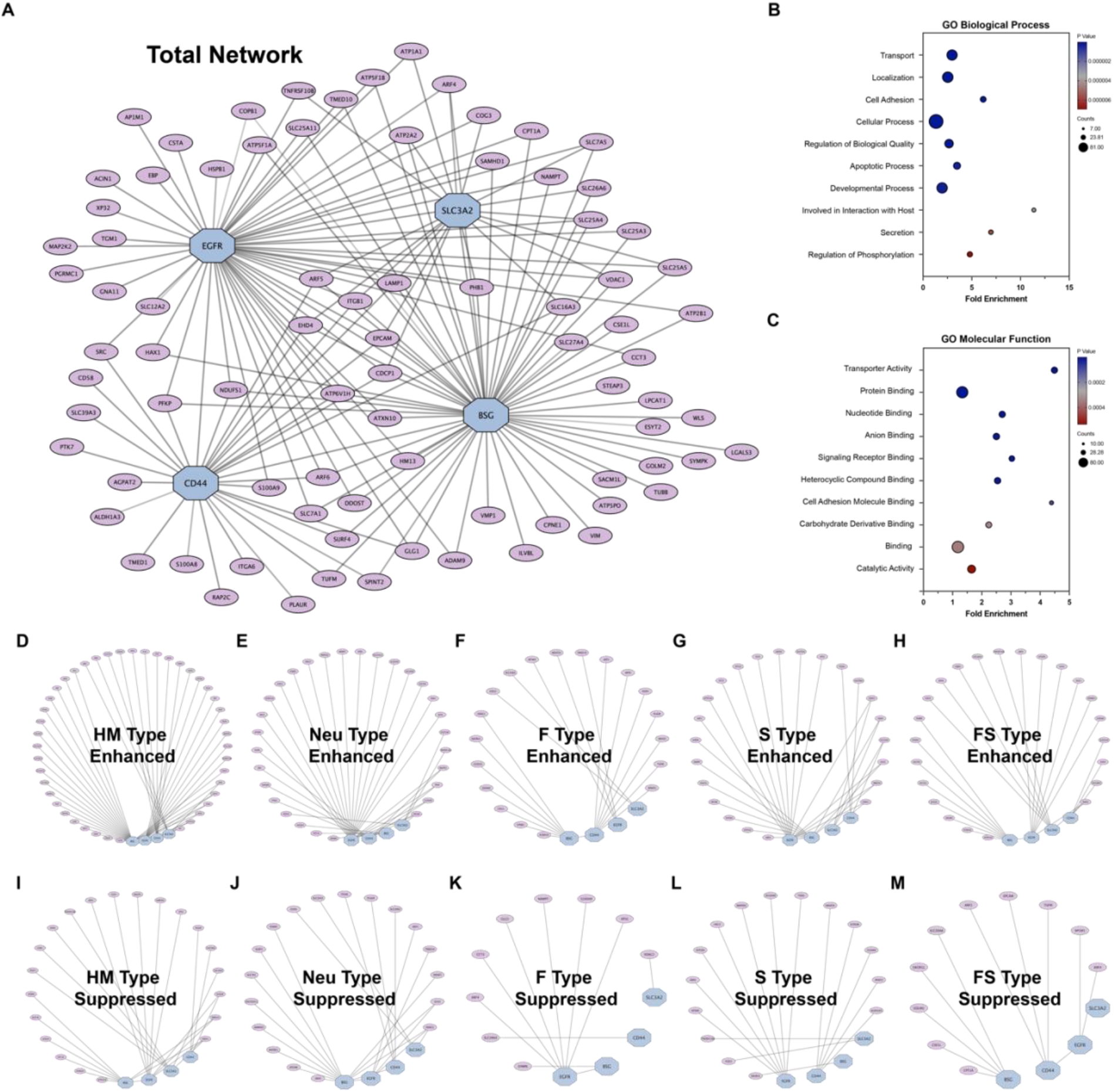
GDPI profile from GAP-MS. **(A)** Overall protein interaction network form AP-MS analysis covering 156 interactions and 85 proteins from four bait glycoproteins, including BASI (Basigin), EGFR (epidermal growth factor receptor), CD44, and SLC3A2 (amino acid transporter heavy chain SLC3A2). **(B)** Annotation of identified proteins by their biological processes revealed an overrepresentation of categories related to transport, localization, cell adhesion, and cellular processes. **(C)** Molecular function analysis identified enriched categories such as transporter activity, catalytic activity, and various binding events. (D)-(H) The enhanced protein subnetworks under HM, Neu, F, S, and FS glycan conditions. (I)-(M) The suppressed protein subnetworks under HM, Neu, F, S, and FS glycan conditions. (All the data is available at www.glycointeractome.org).

In addition, the cell membrane is a highly interactive environment, and many glycoconjugates have been found to form microdomains on the cell surface (Chai et al. 2024). We predicted a high possibility that glycosylated prey proteins could be enriched in our experiments. Therefore, we examined the number of enriched glycoproteins from four baits and found that 26 proteins are glycosylated, spanning 53 interactions. We calculated and compared the glycoprotein enrichment percentage in our experiment to the general membrane proteome, and found the glycosylated protein is indeed more enriched employing our four baits (30% vs 12%). We also counted the interaction edge for different prey proteins, and 47 out of 85 (>55%) were found to have more than one interaction (**Figure S2E**). Interestingly, nearly 60% of the glycoproteins (15 out of 26) were enriched by more than one bait, demonstrating the integrative and complex environment of the cell membrane glycocalyx (**Figure S2F**). Furthermore, clustering proteins based on their biological processes revealed a significant overrepresentation of categories associated with transport, localization, cell adhesion, and cellular process. (**Figure 3B**). As shown in **Figure 3C**, molecular function analysis identified highly enriched categories, including transporter activity, catalytic activity, and different binding events, which closely align with the functions of an active environment on the cell surface. Overall, our results highlight that the GAP-MS workflow facilitates the elucidation of glycoprotein interaction networks.

### Constructing glycan-dependent subnetwork

We then considered the glycan modifier treatments to compare how interactions vary across different glycan phenotypes. Changes in prey protein abundance under various glycan phenotypes were correlated with the effects of glycosylation on specific interaction pairs. For instance, in the EGFR-ACIN1 and BSG-LGALS3 interactions (**Figure S3A**), ACIN1 was more enriched by EGFR in the HM type compared to other types, while LGALS3 was less captured by BSG pulldown in HM. This observation indicates that GAP-MS data can reveal the enhancement and suppression of interactions when the glycan phenotype is altered.

Since samples under different treatments for each replicate were handled in the same batch, we further normalized the quantification values for each interaction pair to the HM type by replicate (**Figure S3B** and **Supplementary Data 3C**) to reduce batch variations. To systematically assess the impact of glycans on the 156 interactions in the network, we applied the topological scoring (TopS) algorithm to these relative quantification results (**Supplementary Data 4A**) (Sardiu et al. 2019). TopS gathers information from the entire input dataset and generates positive and negative scores that reflect the likelihood of whether an interaction pair is true under each glycan phenotype (**Figure S3C**). The wide range of TopS scores provides a clearer indication of whether a certain form of glycosylation plays a positive or negative role in glycoprotein interactions. A larger positive TopS score indicates higher confidence that the interaction pair is enhanced under that glycan phenotype while a more negative score suggests that the interaction is more likely to be suppressed in that type.

In theory, under each glycan phenotype, the 156 interactions in the total network can be categorized into three main groups: (a) interactions boosted by the dominant form of glycans, (b) interactions hampered in that type, and (c) interactions not strongly affected. Based on TopS scores, we generated 5 enhanced subnetworks for all glycan phenotypes, containing interactions with a TopS score > 20, and 5 suppressed subnetworks with interactions having a TopS score < -20 (**Figure 3D-M** and **Supplementary Data 4A**). The 25 interactions that were not included in either the enhanced or suppressed networks in any type were considered independent of the glycan forms we included in the current GAP-MS platform (**Figure S3D and Supplementary Data 4A**). With this cutoff criterion, it is legitimate for an interaction pair to be included in multiple subnetworks as long as it is not in the independent network, such as the example of BSG-LGALS3. From an overview of the subnetworks (**Figure 3D-M**), however, we observed no high similarity between any two subnetworks of different glycan phenotypes. For example, there are 25 interactions in the FS type enhanced network, 20 in the F type enhanced network, and 30 in the S type enhanced network; however, only 4 interactions overlap between the F and FS types, and 11 between the S and FS types.

To reveal any existing patterns of co-occurring glycan dependency within our current dataset, we applied k-means clustering on TopS scores for the 156 interactions under all 5 conditions (**Supplementary Data 4B**). As shown in **Figure 4A**, all interactions in the total network were consistently divided into 6 clusters according to how they were affected by different glycan phenotypes. Examples of interactions from each cluster are displayed in **Figure 4B**, and a summary of each cluster is provided in the table in **Figure 4C**. Clusters 1, 2, and 3 each contain interactions strongly enhanced in one specific glycan phenotype (HM type for Cluster 1, F type for Cluster 2, and Neu type for Cluster 3) while being either moderately suppressed or not notably affected in other types. In Cluster 4, interactions are generally not drastically impacted by any phenotype; the overall pattern suggests they are mostly suppressed in HM and Neu types while mostly enhanced in S and FS types. Cluster 5 contains two distinct modules: interactions are suppressed in HM, Neu, and F types but are strikingly enhanced when sialylation is present (in S and FS types). There are only 5 interactions in Cluster 6, all of which are greatly suppressed in HM type but noticeably enhanced in Neu type.

**Figure 4.**
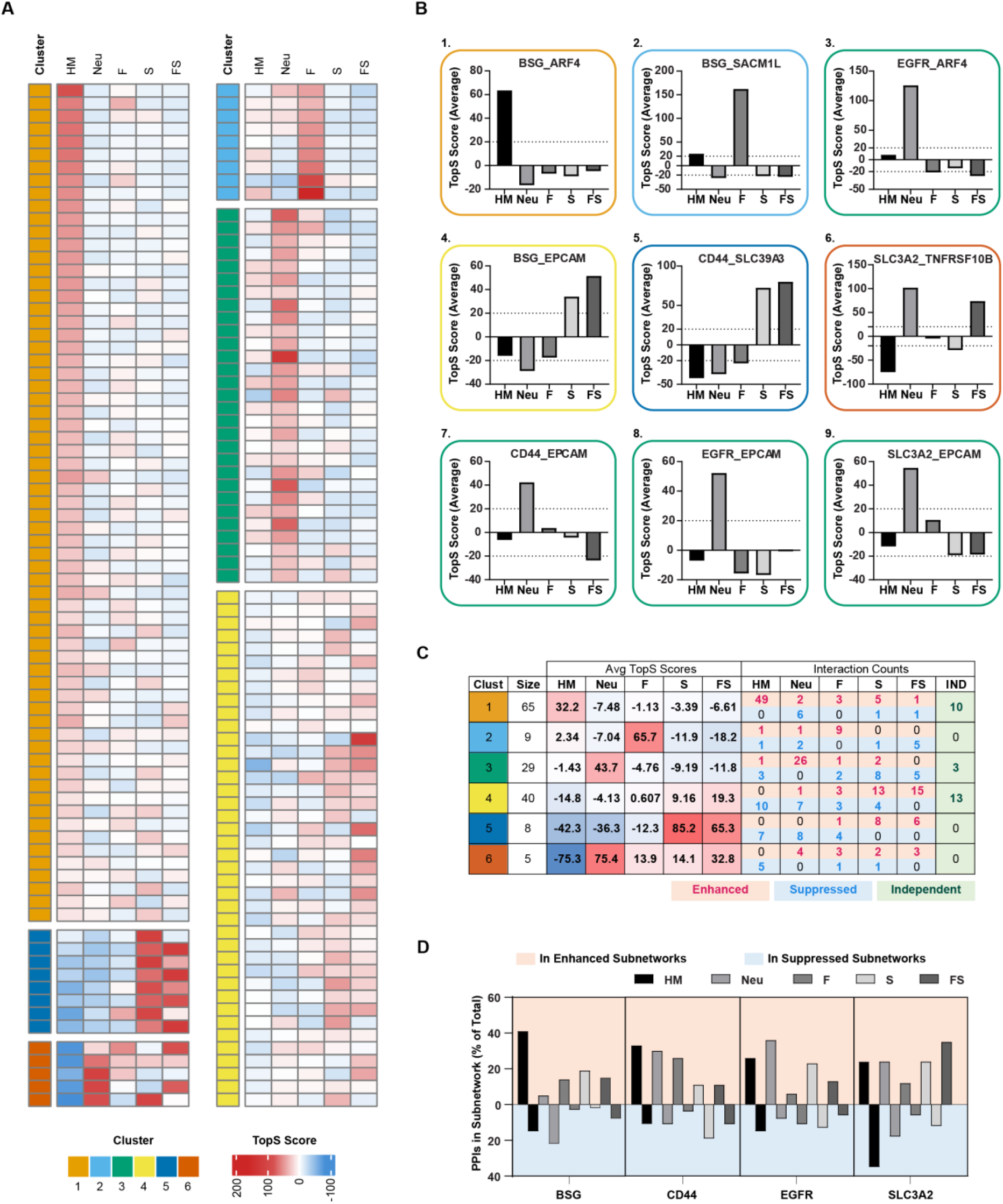
GAP-MS data revealed that different glycan phenotypes have varying effects on interaction pairs. **(A)** Heatmap illustrating the responses of the 156 interaction pairs to each glycan phenotype. A larger positive average TopS score indicates higher confidence that the interaction pair is enhanced, while a more negative TopS score suggests stronger suppression. Based on TopS scores, the 156 interactions were divided into 6 clusters using the k-means method. **(B)** Example interaction pairs from each cluster. **(C)** Summary of interactions within each cluster. **(D)** Summary of interactions in each glycan-dependent subnetwork by bait protein. The percentage was calculated from the number of interactions with each bait in that subnetwork relative to the total number of interactions for the corresponding bait.

As mentioned earlier, glycoproteins on the cell membrane are highly interactive, resulting in some prey proteins being captured by more than one bait. Intriguingly, the interactions of a single prey protein with various bait proteins do not always fall within the same cluster. For example, as shown in **Figure 4B (1)** and **(3)**, the interaction between BSG and ARF4 is in Cluster 1, whereas the interaction between EGFR and ARF4 is in Cluster 3. Another instance involves the prey EPCAM [**Figure 4B (4)** and **(7)-(9)**], which was captured by all four baits. Notably, the BSG-EPCAM interaction is in Cluster 4, while the other three pairs are in Cluster 3. This observation suggests that glycoproteins do not necessarily respond to glycan alterations in the same manner. This is also apparent in **Figure 4D**, where the distribution of pairs in the enhanced or suppressed subnetworks under each glycan phenotype varies for different baits.

### Comparison of GAP-MS with AP-MS combined with glycosite mutagenesis on BSG

Finally, we compared our GAP-MS results with another frequently used approach to study the effect of PTM on protein interactions, wherein we could remove glycosylation from a known site of the bait protein by mutating the asparagine to glutamine. We employed BSG mutagenesis as an example; there are two validated glycosylation sites on BSG, N160 and N268 (corresponding to N44 and N152 in the isoform we expressed). We expressed Halo-tagged BSG containing N160Q, N268Q, or double mutations (DM) in HCT116 cells for AP-MS analysis (**Figure S4A**). One concern with this mutagenesis-based approach is that mutations of certain amino acids in the bait protein could severely compromise its folding and cellular localization. Fortunately, the three forms of BSG with glycosylation site mutations can be normally localized to the membrane like the wild type (**Figure S4B**).

We then evaluated the differences in co-purified proteins of BSG with glycosylation site mutations compared to the wild type (**Supplementary Data 5**). As shown in **Figure S4C-E**, using a limma *p*-value cutoff of 0.05, only 18 proteins were significantly affected by the N160Q mutation, whereas more than 60 proteins were significantly changed in the N268Q and double-mutated BSG pulldown. These results suggest that the second glycosylation site has a greater impact on the protein interactions of BSG. Therefore, we compared the changes in N268Q and double-mutated conditions with the changes in BSG interactions caused by our glycan modifier treatments.

BSG co-purified proteins under different treatments (Neu, F, FS, and HM glycan phenotypes) were compared to the control treatment (S type). Using the same limma *p*-value cutoff of 0.05, significantly changed BSG co-purified proteins in each treatment condition were determined (**Supplementary Data 5**). Shared PPI changes of BSG caused by glycosylation site mutations or global glycan changes are displayed in **Figure S5A**. As also illustrated in the volcano plots (**Figure S4C-E**), the loss of glycosylation sites mainly induces a decrease in many co-purified proteins of BSG. On the other hand, different glycan modifier treatments caused various upregulations or downregulations of interactions. The treatment with Kif or 3-F-Sia mainly enhanced many interactions, while treatment with fucose or both fucose and 3-F-Sia caused a similar number of increased and decreased interactions. Most increased or decreased interactions caused by the N268Q and double mutations overlapped. However, it’s worth noting that very few changes were shared between the mutagenesis conditions and any of the glycan modifier treatments. These results demonstrate that our GAP-MS workflow provides distinct insights compared to those obtained from the mutagenesis approach.

## DISCUSSION

In this study, we introduced the GAP-MS system, designed to systematically investigate the glycoprotein interactome using AP-MS under five conditions with controlled alterations of global glycosylations (**Figure 1A-D**). This novel AP-MS-based workflow enabled the uncovering of GDPI for any glycoprotein of interest (**Figure 1E**), providing information that has been difficult to obtain with existing approaches. The critical feature of GAP-MS lies in the integration of metabolic manipulation of glycan types in cell culture (**Figure 1B-C**). Treatments with one or a combination of glycan modifiers for a sufficient period have been tested to consistently change the glycan profiles (**Figure 2A**). We introduced the concept of glycan phenotype to describe these outcome changes in cells. The glycan modifiers can be directly fed to cells, thus avoiding the introduction of genetic variations when comparing different glycan phenotypes. This manipulation approach, combined with DIA-based high-resolution mass spectrometry analysis, allows GAP-MS data collected at different time points to be aggregated later. Beginning with this study, the GDPI-derived networks can continue to expand in the future as more glycoproteins of interest are analyzed as baits and additional glycan phenotypes are generated.

Using BSG, CD44, EGFR and SLC3A2 as baits to prove the principle, we identified 156 interactions with high confidence and illustrated how they were influenced by the five glycan phenotypes involved in this study (**Figure 3**). In each type, interactions were either enhanced or suppressed (**Figure 3D-M**). This key observation clearly shows that different glycan forms play distinct roles in different proteins or interactions. This is even more evident when examining glycan-dependent interactions by bait (**Figure 4D**). For example, the HM type predominantly enhances interactions of BSG, CD44, and EGFR, while it plays a negative role in most interactions of SLC3A2. Increased protein binding mediated by high-mannose glycosylation has been reported to play critical biological roles (Heller et al. 2003; Park et al. 2020). Our data suggest that an overabundance of high-mannose glycans may also lead to the loss of certain protein interactions, potentially contributing to the molecular and cellular changes associated with dysregulated high-mannose levels. This observation is not unique to the HM type, similarly, none of the other types of glycosylations exhibit the same effects across all baits. This can also be seen in some of the interaction examples shown in **Figure 4B**, where interactions with the same prey protein respond differently to glycan phenotype changes depending on the bait. Since the manipulation of glycans in GAP-MS affects all glycoproteins that undergo glycosylation after the treatments, it becomes challenging to determine whether the glycans on the bait or the prey protein—or even the specific glycosylation site—play the more pivotal role. In such cases, other approaches may serve as useful complements to further investigate the specific interactions identified by GAP-MS. For instance, we applied mutagenesis to the glycosylation sites of BSG and performed AP-MS analysis (**Figure S4**). This data clearly indicates that the loss of the N268 site has a more pronounced impact on interactions than the loss of the N160 site. For example, the loss of only N160 barely affected the pulldown of LGALS3, whereas the additional loss of N268 caused a substantial reduction in the same prey (**Figure S5C**). There are also interactions affected by both sites; for instance, the loss of either N160 or N268 significantly increased the pulldown of CDCP1, and losing both resulted in an even more significant increase (**Figure S5D**). Despite this limitation, GAP-MS provides unique insights. Taking the interaction between BSG and CDCP1 as an example, GAP-MS reveals that the HM type significantly increased the co-purification of CDCP1 with BSG (**Figure S5E**), suggesting that the glycosylation of BSG does not always inhibit this interaction.

In GAP-MS, for each pair of confident interactions, the influence of various glycan phenotypes is also illustrated in parallel (**Figure 4A**). This allows for observing how different glycans may employ similar or opposing effects on the same interaction pair. The clustering analysis highlights groups of interactions regulated by glycans similarly (**Figure 4C**). Among the 156 interactions across the 5 glycan phenotypes we examined, we found that HM, F, and Neu each play a major role in enhancing interactions within their corresponding clusters (Clusters 1, 2, and 3). For interactions in these three clusters, the other glycan types do not exhibit drastic effects. Interactions in Cluster 4 are not strongly impacted by any single glycan phenotype, though there is a slight boost for glycans with fucose or sialic acid units (F, S, and FS) compared to HM and Neu. Intriguingly, only small groups of interactions seem to be predominantly influenced by the presence of a single type of monosaccharide (Clusters 5 and 6). In Cluster 5, interactions are strongly enhanced by sialylation (present in S and FS), while in Cluster 6, interactions are clearly dependent on the presence of galactose, leading to strong suppression in HM. However, only 13 interactions from these two clusters are found out of a total of 156. In other words, GAP-MS results suggest that, in most cases, it is the overall glycan structure that determines how glycosylation affects protein interactions.

The application of controlling glycan phenotypes is also versatile. In this work, we combined it with AP-MS for robust and confident identification of PPIs, but it can also be integrated with other methods such as proximity-based labeling and crosslinking mentioned earlier. Due to the nature of affinity purification, interactions mediated either directly or indirectly by glycans are all captured. For proteins that bind to glycans or regions near glycosylation sites, it is reasonable to expect that their interactions will be affected when glycan structures change. Taking the BSG-LGALS3 pair as an example (**Figure S3A-C**), this interaction is strongly downregulated in the HM type, while the other four types enhance it. This is due to the lack of galactose in the HM type, which is consistent with the fact that LGALS3 is a galactose-specific lectin (Joeh et al. 2020). On the other hand, proteins that interact indirectly with glycoproteins or bind far from the glycosylation sites can also be captured by AP-MS. It is also reasonable to expect that these interactions may not be affected by changes in glycan phenotypes. This explains the interactions in **Figure S3D**, which we classified as independent of glycans. However, one of the most intriguing aspects of GAP-MS is that some of these interactions, which are not expected to be influenced by glycans, are found to be glycan-dependent. One example is shown in **Figure 4B(3)**, where the interaction between EGFR and ARF4 is strongly enhanced in the Neu type. It is known that ARF4 binds to the cytoplasmic domain of EGFR, while the glycosylation occurs on the extracellular part of EGFR (Kim et al. 2003). This might be explained by conformational changes induced by glycosylation, which could be further investigated using the glycan modifiers with other structural analysis approaches in future studies.

## RESOURCE AVAILABILITY

### Lead contact

Further information and requests for resources and reagents should be directed to and will be fulfilled by the lead contact, Yixuan Xie.

### Materials availability

All the MS raw data generated in this study have been deposited to the Mass Spectrometry Interactive Virtual Environment (MassIVE) repository with the dataset identifier MSV000096043

## Data and code availability

All the interactome results have been deposited at: www.glycointeractome.org.

## ACKNOWLEDGMENTS

This work was supported by grants from the Greater Bay Area Institute of Precision Medicine (Guangzhou) I0036(A) (Y.X.), the National Institutes of Health GM049077 (C.B.L.), AG062240 (C.B.L.), AI118891 (B.A.G.), HD106051 (B.A.G.), and CA196539 (B.A.G.). Z.L. was supported by the Research Education Component (REC) through the NIA P30 AG066444 grant. The authors thank the suggestions for using TopS and LIMMA from Dr. Mihaela E. Sardiu at the University of Kansas. The authors also thank the support received from Dr. Wandy Beatty and the Molecular Microbiology Imaging Facility at Washington University School of Medicine.

## AUTHOR CONTRIBUTIONS

X.L. and Y.X. convinced the project. X.L., B.A.G., and Y.X. designed the overall plan. X.L., L.Y., S.C., S.Y., and Y.X. performed the experiments. X.L., L.Y., Z.L., S.W., and Y.X., analyzed data. X.L., L.Y., and Y.X. produced figures. X.L. and Y.X. drafted the manuscript. L.Y., Z.L., S.C., S.W., C.B.L., and B.A.G. reviewed and edited the manuscript. C.B.L., B.A.G., and Y.X. supervised the overall project.

## EXPERIMENTAL MODEL AND STUDY PARTICIPANT DETAILS

### Mammalian cell culture

HCT116 cells were maintained in Dulbecco’s Modified Eagle Medium (DMEM) supplemented with 10% Fetal Bovine Serum (FBS), MEM nonessential amino acids (NEAA) and GlutaMAX™.

## METHOD DETAILS

### Fluorescent imaging

Wildtype HCT116 cells or cells stably expressing Halo tagged proteins were plated in a 35-mm MatTek dish with No. 1.5 glass bottom coated by poly-d-lysine. For transient expression, transfection was performed on the next day after plating cells. Cells expressing Halo proteins were stained with HaloTag® TMRDirect™ Ligand (Promega, Madison, WI, USA) overnight in their regular culture medium. On imaging day, cells were stained with CellMask™ Green Plasma Membrane Stain (Invitrogen, Carlsbad, CA, USA) and Hoechst33342 for 15 minutes in a 37 **°C** incubator. For the stable cell lines, stained samples were fixed with 4% formaldehyde and washed with PBS. Fixed samples were imaged in VECTASHIELD® Antifade Mounting Medium (Vector Laboratories, Newark, CA, USA). For transiently transfected samples, cells were washed with a warm culture medium after staining and imaged as live cells in FluoroBrite™ DMEM (Invitrogen, Carlsbad, CA, USA) supplied with 10% FBS. Imaging of both fixed and live cell samples was performed on a Zeiss LSM 880 Confocal Laser Scanning Microscope (Carl Zeiss Inc. Thornwood, NY, USA). The base of the microscope is inverted and cells were imaged through a Plan-Apochromat 40x/1.4 Oil objective. HaloTag® TMRDirect™ was excited by 543nm helium-neon laser and detected at 553-753 nm. Hoechst33342 was excited by 405nm diode laser and detected at 415-470 nm. CellMask™ Green was excited by 488nm argon laser and detected at 491-553 nm. ZEN black software (version 2.1 SP3) was used for multichannel image acquisition and analyses. Single color control experiments were performed separately for HaloTag® TMRDirect™ and CellMask™ Green to make sure no crosstalk between channels.

### Affinity purification

Each cell line stably expressing Halo only or Halo-tagged bait protein was plated at a density of 2 million cells per 100mm plate in medium without Hygromycin B. Glycan modifiers were fed to cells at working concentration on the next day and cells were collected 3 days after treatment. Halo purification was performed according to the manual of HaloTag® Mammalian Pull-Down Systems (Promega, Madison, WI, USA) with minor optimizations. In detail, each cell pellet from the 100-mm plate was lysed with the Mammalian Lysis Buffer by passing by a 28 gauge needle 5 times. Crude lysates were centrifuged at 10,000 × *g* at 4 °C for 10 min, supernatants were collected to bind with Magne® HaloTag® Beads. After rotating at 4°C overnight, beads were washed with cold Wash Buffer 5 times then eluted with AcTEV Proteoase (Invitrogen, Carlsbad, CA, USA) at room temperature for 1 hour with shaking. Eluats were subjected to sample processing for mass spectrometry analysis. For transient expression, HCT116 cells were plated in the medium without any antibiotics. Cells were collected 48 hours after transfection, and halo purification was performed the same as described above.

### Glycomic analysis

The glycomic samples were prepared as described previously (Li et al. 2020). Briefly, the enriched membrane was resuspended with 200 µL of 100 mM HEPES buffer, and the mixture was heated using a thermomixer at 100 °C for 2 min. The *N*-glycans cleavage was performed by adding 2 µL of PNGase F (500,000 units/mL), followed by incubation at 37 °C overnight. The supernatant containing the released *N*-glycans was purified using the PGC plate using the porous graphitic carbon (PGC) SPE plate (Thermo Scientific, MA) and was eluted with 60% (*v*/*v*) ACN and 0.1% (*v*/*v*) TFA in water. The purified glycans were dried using the SpeedVac system (Thermo Scientific, MA) and reconstituted in water. The sample was analyzed using a 1200 Series liquid chromatography chip system (Agilent, CA) coupled with a 6520 Accurate Mass Q-TOF system (Agilent, CA). The glycans separation was carried out at a constant flow rate of 0.3 μL/min using buffer A (water containing 0.1% formic acid) and buffer B (acetonitrile containing 0.1% formic acid). The chromatography gradient consists of 0−2 min, 0% B; 2−20 min, 0-16% B; 20−40 min, 16%-72% B; 40−42 min, 72%-100% B; 42-52 min, 100% B; 52-54 min 100%-0% B; 54-65 min 0% B.

The glycans were identified using MassHunter Qualitative Analysis B. 07 software (Agilent, CA). Relative abundances of each *N*-glycan subtype were calculated after normalizing the integrated peak areas to the total peak areas of all glycans detected.

### Glycoproteomic analysis

The glycopeptides were enriched by solid-phase extraction using iSPE®-HILIC cartridges (The Nest Group, MA) after the tryptic digestion, and the samples were analyzed using a Vanquish Neo UHPLC System coupled with an Orbitrap Exploris 240 mass spectrometer. 2 μL of the sample was injected, and the analytes were separated on an EASY-Spray PepMap Neo Column (3 μm, 0.075 mm × 150 mm, Thermo Scientific, CA). LC separation was performed with a binary gradient using solvent A with 0.1% (*v*/*v*) formic acid (FA) in water and solvent B with 0.1% (*v*/*v*) FA in ACN at a flow rate of 300 nL/min. After the separation, the peptides were analyzed with the full MS scan from 700 to 2000 in positive ionization mode. The MS/MS spectra were collected for fragments with m/z values starting from 120. Glycopeptides were identified using Byonic software (Protein Metrics, CA). Raw files were searched against a human protein FASTA database acquired from UniProt. C-Terminals of lysine and arginine were used for specific cleavage sites, and missed cleavages were restricted to 2. Precursor mass tolerance was limited to 10 ppm, and CID & HCD fragmentation with a mass tolerance of 20 ppm was applied. Carbamidomethylation at cysteine was assigned as the fixed modification. An in-house human database was applied for N-glycosylation at asparagine, and identifications with high confidence were achieved through filtering with a 1% false discovery rate (FDR), score greater than 300, and DeltaMod larger than 10.

### Proteomic analysis

The lysis buffer consisting of 5% SDS in 50 mM triethylammonium bicarbonate buffer (TEAB, Sigma-Aldrich, MO) was added to the cell pellet, and the cells were fully lysed using sonication at room temperature for 5 min. Reduction and alkylation of proteins were performed by adding 1 μL of 200 mM tris(2-carboxyethyl)phosphine hydrochloride (TCEP, Sigma-Aldrich, MO) at 55 °C for 15 min and 1 μL of 400 mM iodoacetamide (IAA, Sigma-Aldrich, MO) at room temperature for 20 min. The proteins were further loaded to the S-trap micro columns (ProtiFi, NY) and cleaned based on the manufacturer protocols. The proteins were digested with 2 μg of trypsin (Promega, WI) at 37 °C overnight. The digested peptides were eluted using 50 mM ammonium bicarbonate buffer, 0.2% formic acid (FA, Fisher Scientific, NH), and 50% acetonitrile (ACN, Thermo Scientific, MA), respectively. Tryptic digested samples were reconstituted in 0.1% FA and characterized using a Vanquish Neo UHPLC System (Thermo Scientific, CA) coupled with an Orbitrap Exploris 240 mass spectrometer (Thermo Scientific, CA). The analytes were separated on an EASY-Spray PepMap column (3 μm, 0.075 mm × 150 mm, Thermo Scientific, CA) at a flow rate of 0.3 μL/min, and the column temperature was set at 45 °C. A solution of water containing 0.1% FA and acetonitrile containing 0.1% FA were used as solvents A and B, respectively. The chromatography gradient consisted of 2% solvent B over 0-2 min, 2-32% solvent B over 2-75 min, 32-45% solvent B over 75-80 min, 45-80% solvent B over 1 min, 80% solvent B over 81-86 min, and finally equilibrated with 2% solvent B over 4 min. The Orbitrap Exploris 240 was equipped with an EASY-Spray Source (Thermo Scientific, CA). The DIA method consisted of staggered DIA windows that spanned the mass range *m/z* 400-1000, and the acquisition settings were as follows: MS1 accumulation time 60 ms, MS2 accumulation time 60 ms, and MS2 first mass at *m/z* 120 . All raw data were processed with Spectronaut (v18.3) using the directDIA mode supplied with the additional library. The peptides were searched using the following parameters: Trypsin/P, one missed cleavage allowed, N-term M excision, fixed modification: C carbamidomethylation, variable modification: none, peptide length range: 7 to 30 amino acids.

## QUANTIFICATION AND STATISTICAL ANALYSIS

The identified protein was subjected to protein-protein interaction scoring with SAINTexpress with default settings (Teo et al. 2014). To generate the overall interaction network, we included co-purified proteins with a SAINT score ≥ 0.90 for each bait. The different glycan topological networks were further established based on TopS scores.(Sardiu et al. 2019) For input to TopS, raw protein quantifications exported from Spectronaut were first normalized to the bait protein and then transformed into relative abundances compared to the high-mannose type condition. Interactions with TopS scores above 20 or below -20 were included to generate the glycan-dependent subnetworks for each glycan phenotype. The total interaction network and all subnetworks were visualized using Cytoscape (v.3.9.1) (Shannon et al. 2003). For k-means clustering analysis, the number of clusters (k = 6) was determined using the silhouette method with the *R* package factoextra (Kassambara 2016). With the nstart parameter set to 25, the same clustering result was consistently reproducible. LIMMA analysis was performed with StatsPro, using the Spectronaut-reported quantifications as input (van Ooijen et al. 2018; Yang et al. 2022). For comparison of BSG AP-MS data between conditions, a LIMMA p-value of 0.05 was used to determine significant differences. Only significantly altered co-purified proteins for each mutation or glycan modifier treatment were included in the overlap analysis. UpSet plots were generated in *R* using the ComplexHeatmap package in intersect mode(Gu, Eils, and Schlesner 2016; Gu 2022).

---

**Figure S1.**
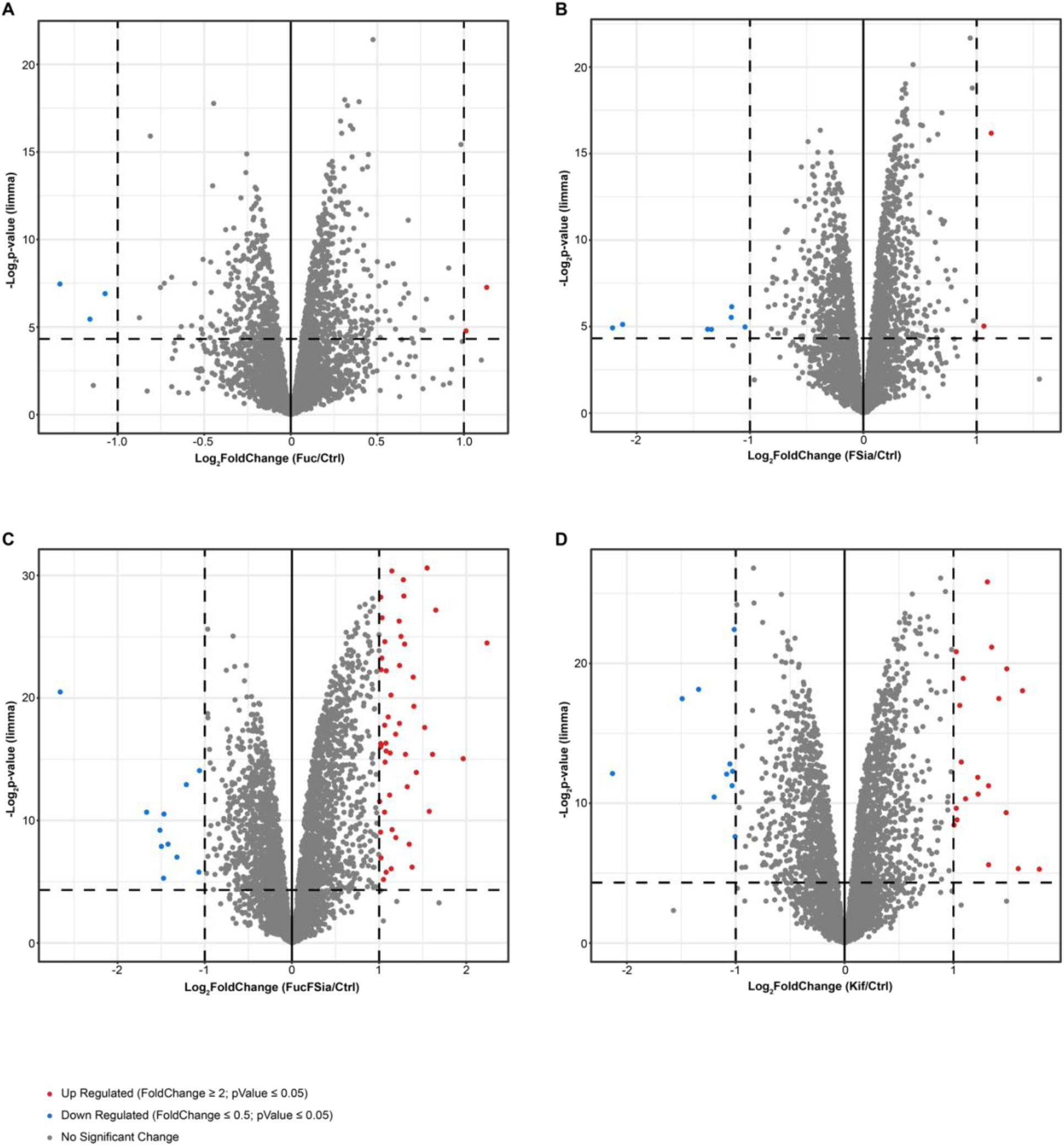
Volcano plots displaying changes in whole proteome abundance following treatment with glycan modifiers. Changes with p-values ≤ 0.05 and an absolute fold change greater than 2 are considered significant and are highlighted in color in the plots.

**Figure S2.**
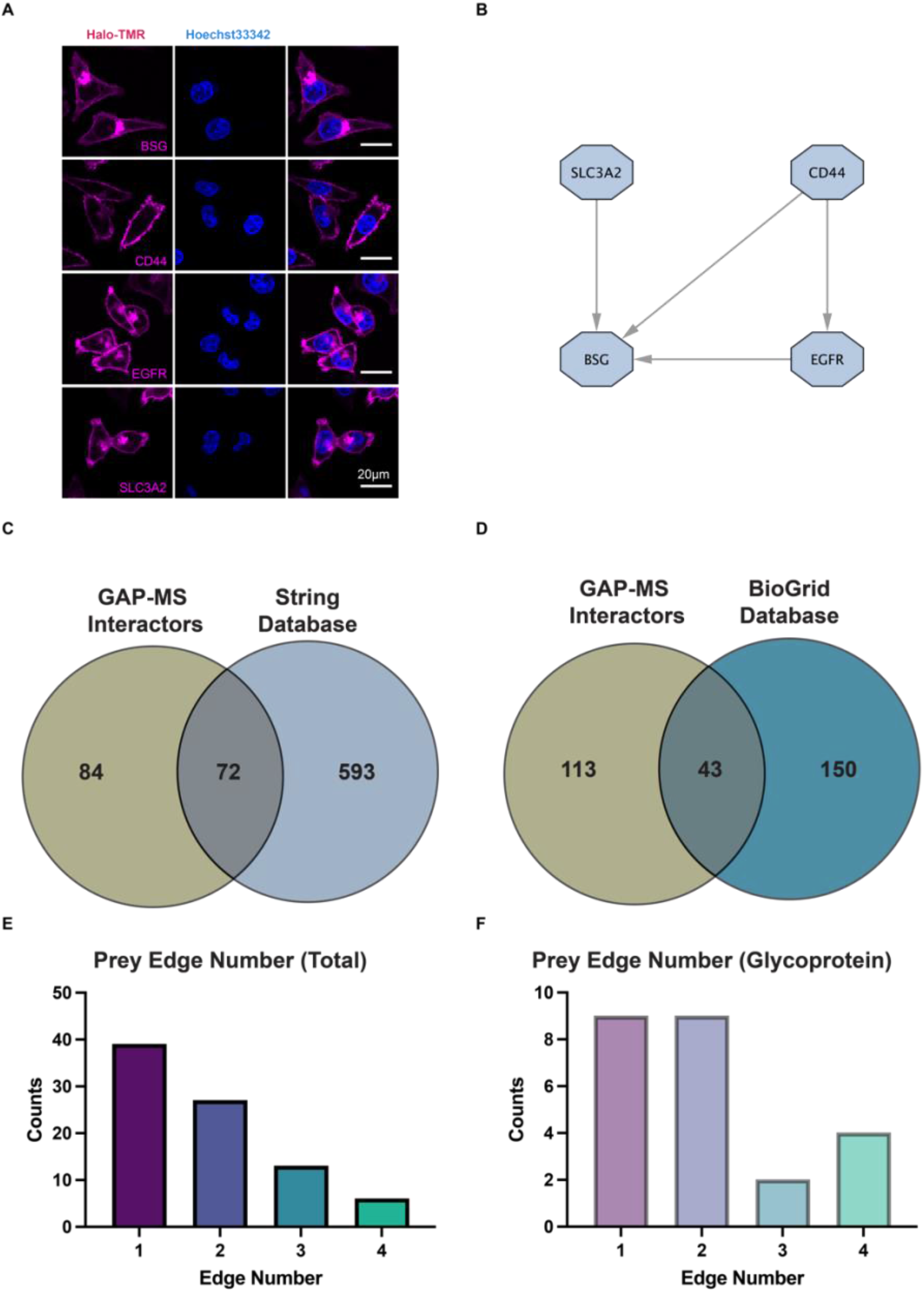
Supplementary figures for performing GAP-MS analysis on selected glycoproteins using HCT116 cells stably expressing Halo-tagged baits. **(A)** Fluorescent confocal microscopy images of HaloTag in cell lines stably expressing each Halo-tagged bait protein. Bait proteins are shown in magenta (pseudo color) and nuclear staining is shown in blue (pseudo color). All scale bars represent 20 microns. All four Halo-tagged bait proteins (BSG, CD44, EGFR, and SLC3A2) marked the cell outlines, indicating that the tag and overexpression did not disrupt the localization of these glycoproteins to the cell membrane. Halo-tagged BSG, EGFR, and SLC3A2 also displayed puncta inside the cells, not overlapping with the nucleus, which may represent accumulation of the overexpressed proteins in the Golgi. **(B)** The extracted module from the total PPI network containing only the bait proteins. **(C)** and (D) GAP-MS results compared to String and BioGrid Databases, respectively. (E) and (F) Summary of egde counts for total and glycosylated prey proteins and prey proteins.

**Figure S3.**
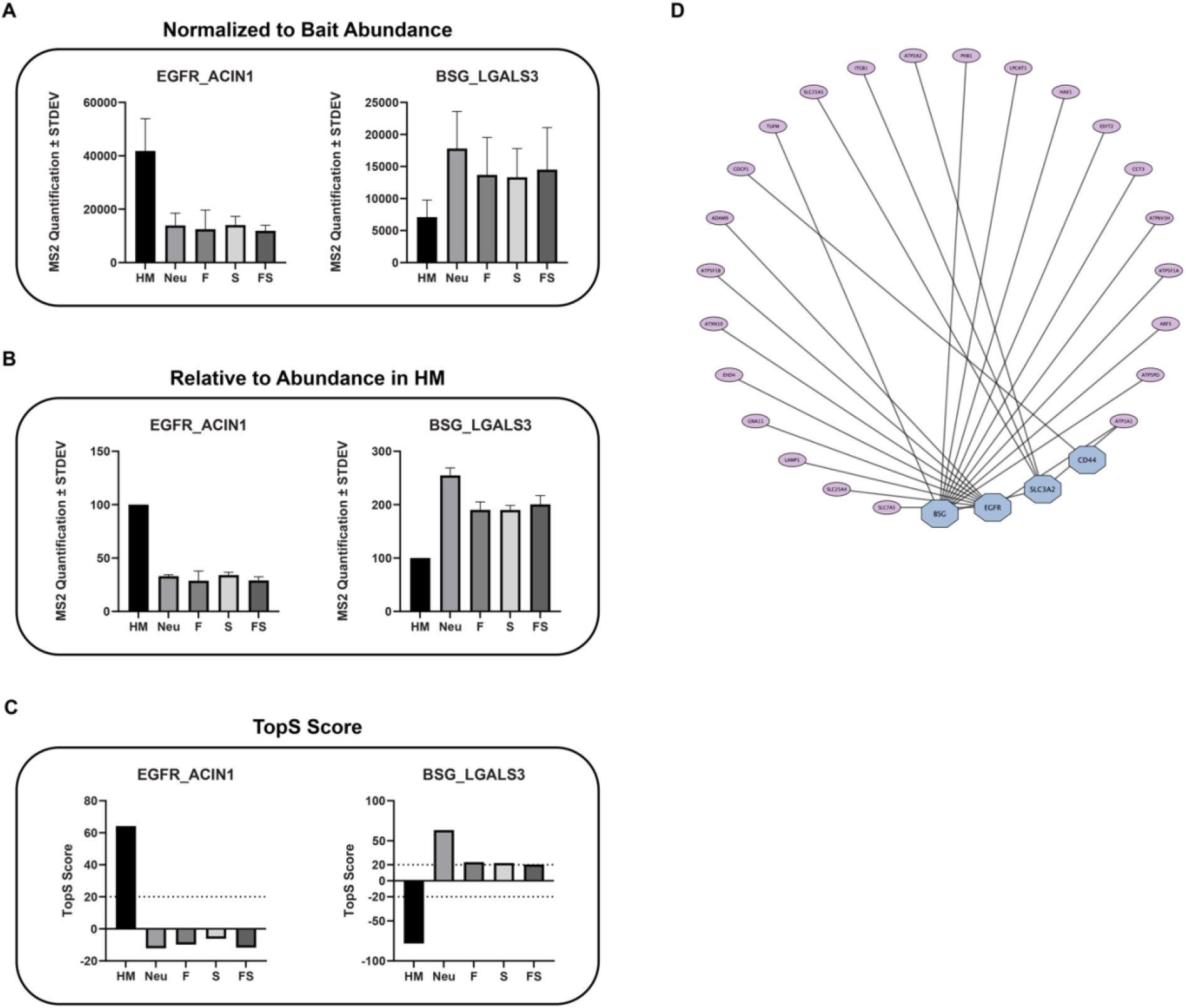
Supplymentary figures demonstrating the use of GAP-MS data to reveal glycan-dependent PPI networks. (A)-(C) Different stages of data processing from Spectronaut-reported MS2 quantifications, illustrated with two example interaction pairs: EGFR-ACIN1 and BSG-LGALS3. In panel A, the abundance of the co-purified prey in each type was normalized by the bait abundance in the corresponding sample. In panel B, the normalized abundance in the HM type was set to 100, and the other types within the same batch (replicate) were transformed to relative abundance compared to the HM type. Finally, these relative abundances were used as input to compute the average TopS scores, shown in panel C. (D) The glycan-independent interactions identified by the current GAP-MS data.

**Figure S4.**
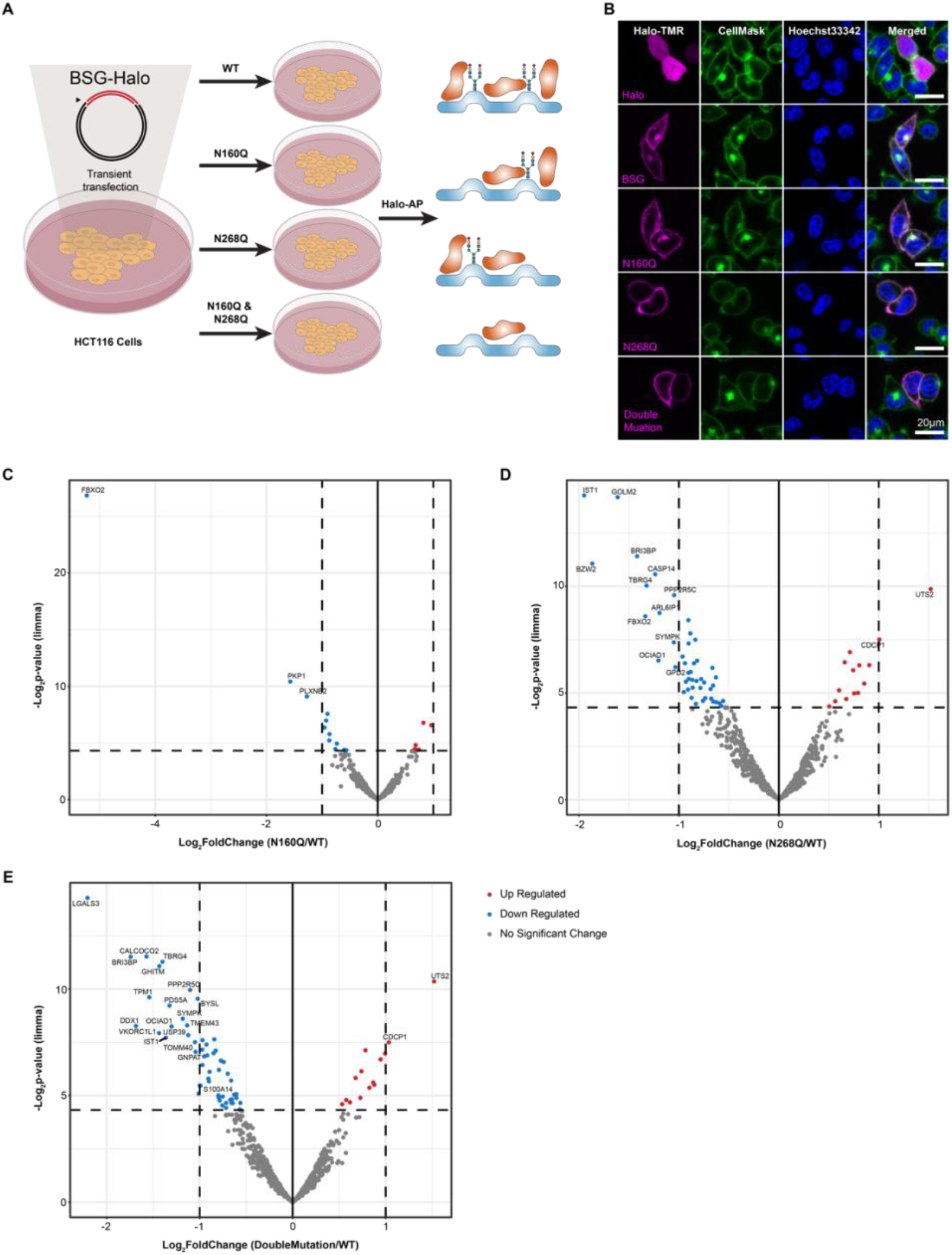
Glycosite mutagenesis on BSG followed by AP-MS analysis. **(A)** Schematic overview of the workflow. **(B)** Live-cell fluorescent imaging of HCT116 cells transiently expressing Halo-tagged wild-type or mutated BSG, or HaloTag alone. Each Halo-tagged protein or the tag alone is shown in magenta (pseudo color), the CellMask reagent staining the cell membrane is shown in green (pseudo color), and nuclear staining is shown in blue (pseudo color). All scale bars represent 20 microns. The images show that wild-type and glycosylation site-mutated versions of BSG localize to the cell membrane as expected, while HaloTag alone is distributed throughout the cell. (C)-(E) Volcano plots displaying changes in co-purified proteins with BSG when its glycosylation sites were removed by mutagenesis. Changes with p-values ≤ 0.05 are considered significant and are highlighted in color in the plots. Proteins with an absolute fold change greater than 2 are labeled.

**Figure S5.**
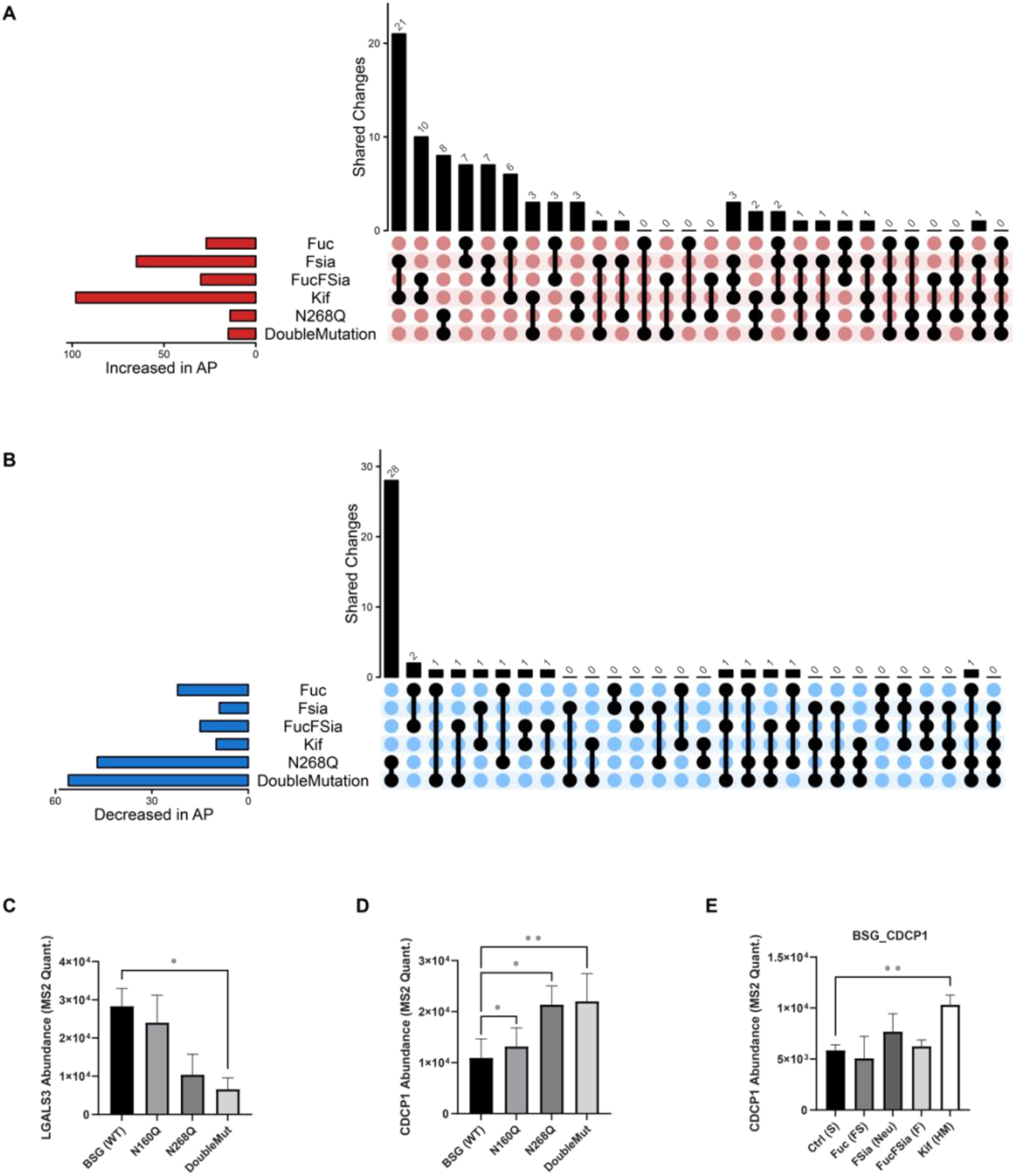
Comparison of the GAP-MS approach to glycosite mutagenesis followed by AP-MS. (A) and (B) UpSet plots illustrating the overlaps of significantly changed co-purified proteins with BSG under each glycan modifier treatment or with glycosylation site mutations on the bait protein. Panel A compares the significantly increased preys, while panel B compares the significantly decreased preys. (C)-(E) Additional examples of interactions shown in unprocessed MS2 quantifications presented as bar plots. In addition to the limma method (results included in Supplementary Data 5), paired t-tests were also performed for these examples. Two-tailed p-values were used, and significant differences between conditions are marked in the plots.

